# Structure-selected RBM immunogens prime polyclonal memory responses that neutralize SARS-CoV-2 variants of concern

**DOI:** 10.1101/2021.10.01.462840

**Authors:** Gonzalo Almanza, Valentina Kouznetsova, Alex E. Clark, Eduardo Olmedillas, Andrea Castro, Igor F. Tsigelny, Yan Wu, George F. Gao, Erica Ollmann Saphire, Aaron F. Carlin, Maurizio Zanetti

## Abstract

Successful control of the COVID-19 pandemic depends on vaccines that prevent transmission. The full-length Spike protein is highly immunogenic but the majority of antibodies do not target the virus: ACE2 interface. In an effort to concentrate the antibody response to the receptor-binding motif (RBM) we generated a series of conformationally-constrained immunogens by inserting solvent-exposed RBM amino acid residues into hypervariable loops of an immunoglobulin molecule. Priming C57BL/6 mice with plasmid (p)DNA encoding these constructs yielded a rapid memory response to booster immunization with recombinant Spike protein. Immune sera antibodies bound strongly to the purified receptor-binding domain (RBD) and Spike proteins. pDNA primed for a consistent response with antibodies efficient at neutralizing authentic WA1 virus and two variants of concern (VOC), B.1.351 and B.1.617.2. These findings demonstrate that immunogens built on structure selection can focus the response to conserved sites of vulnerability shared between wildtype virus and VOCs and induce neutralizing antibodies across variants.

## Introduction

The current SARS-CoV-2 pandemic has been globally disruptive (Nicola *et al*, 2020). Nonpharmaceutical interventions (NPI) have reduced viral transmission but are difficult to sustain due to deleterious social and economic consequences [https://data.undp.org]. Additionally, NPI, as implemented, have not sufficiently controlled a global pandemic that has caused more than 219 million infections and 4.55 million deaths (https://coronavirus.jhu.edu/map.html). Therefore, a systematic deployment of vaccines is required to generate protective immunity and contain virus transmission.

The most effective way to control infectious agents at the population level is immunization and virtually all licensed vaccines owe their protective effects to the induction of pathogen-specific antibodies (Plotkin, 2010). Vaccine-induced antibodies protect either by preventing infection, i.e., blocking the interaction of a virus with its cell target (e.g., lung cells in the case influenza virus), or by preventing disease, i.e., blocking the virus from reaching its target organ (e.g., the central nervous system in the case of paralytic poliovirus). Cellular immunity by T cells and NK cells protect from pathology and disease by killing virus-infected cells (Sridhar *et al*, 2013) or, more generally, by limiting harmful consequences of immune activation (de Jong *et al*, 2006). Therefore, community spread of infection is preferably controlled by antibodies that intercept virions preventing them from binding their receptor on target cells.

Every protein immunogen is composed of various B cell and T cell epitopes against which the immune system responds using its adaptive arm. Polyclonal antibody responses are by definition heterogeneous, are driven by inter-clonal competition (Altman *et al*, 2018; Kuraoka *et al*, 2016), and favor the response to some epitopes at the expense of others, a phenomenon known as immunodominance (Cirelli *et al*, 2019). As a consequence, not all epitopes in a viral pathogen induce responses beneficial to the host. For example, some antigens (e.g., nucleocapsid protein) are immunogenic and have diagnostic value (Burbelo *et al*, 2020) but the immune response against them will not prevent infection. Other epitopes suppress the immune response (Xu *et al*, 2018), or may induce antibodies that exacerbates pathogenesis (Arvin *et al*, 2020). To minimize immunodominance by irrelevant B cell epitopes and their negative impact on the immune response (Altman *et al*, 2018), the immune response should be controlled by narrowing the choice of B cell epitopes involved to those with the highest probability of inducing antibodies against sites of vulnerability on the virus.

The receptor binding motif (RBM) ridgeline contributes numerous amino acid residues to the interaction with ACE2 (Lan *et al*, 2020; Starr *et al*, 2020), it is the target of potent neutralizing antibodies isolated from convalescent individuals via VH/VL cloning (Barnes *et al*, 2020b; Liu *et al*, 2020; Pinto *et al*, 2020; Rogers *et al*, 2020; Shi *et al*, 2020; Tortorici *et al*, 2020; Wu *et al*, 2020; Zost *et al*, 2020) even though the > 80% of the whole antibody response to the Spike protein in convalescent individuals is directed predominantly to sites outside the receptor binding domain (RBD) (Voss *et al*, 2021). Furthermore, patients with mild disease and those with severe disease generate antibodies that tend to recognize different sites in the RBD (Ravichandran *et al*, 2021). Residues in the RBM involved in ACE2 contact are necessarily constrained (Starr *et al*., 2020) with only two common amino acid mutations in SARS-CoV-2 variants of concern (VOC) in the RBM ridgeline region (E484K in B.1.351, P.1 and a small minority of B.1.1.7 and T478K in B.1.617.2). Although neutralizing antibodies have been mapped to the N-terminal domain (Chi *et al*, 2020; Voss *et al.*, 2021) or other sites distal from the RBM (Lv *et al*, 2020; Pinto *et al.*, 2020; Yuan *et al*, 2020), all but one of the 20 most potent (IC50 < 0.1 μg/mL) characterized to date neutralizing antibodies bind the RBM and block receptor attachment (Dejnirattisai *et al*, 2021).

Here we used protein engineering to generate three plasmid (p)DNA immunogens expressing a B cell epitope of the RBM ridgeline, all comprising the ^486^Phe-Asn-Cys-Tyr^489^ “FNCY” patch. This patch plays a key role at the RBM: ACE2 interface, is solvent-exposed, and is included in the epitope target of some of the most potent human Class 1 neutralizing antibodies (Barnes *et al*, 2020a; Dejnirattisai *et al*., 2021). pDNA priming of C57BL/6 mice induced a primary and memory antibody responses against RBD. Antibodies expanded by booster immunization were highly effective at neutralizing authentic WA1 virus and the B.1.351 and B.1.617.2 VOCs. This approach shows that it is possible to initiate anti-SARS-CoV-2 responses recruiting B cell with receptors complementary to a narrow region of the RBM.

## Results

### Prime-boost immunization and serum antibody response in mice

We utilized protein engineering to express RBM amino acid sequences in the complementarity determining regions (CDRs) of the variable (V) domain of an immunoglobulin (Ig) molecule, antibody antigenization (Zanetti, 1992). CDRs are solvent-exposed hypervariable loops supported by the conserved Ig fold (Wu & Kabat, 1970) and independent of the physicochemical constraints that maintain the VH and VL packing. CDRs are ideal sites to express heterologous B-cell epitopes with constrained geometry imparting them with antigenicity and immunogenicity (Zanetti & Billetta, 1996).

An analysis of the RBD: ACE2 interface revealed multiple contact points involving conserved residues on both sides (Fig.1A). We selected a putative B cell epitope comprising a solvent-exposed patch of four amino acid (FNCY) as a target of B cell immunity (Fig. 1B) as several human Class 1 neutralizing antibodies (Barnes *et al*., 2020a) have paratopes that target this site on the RBM ridgeline (Fig. 1C). We generated three immunogens (referred hereunder as Model 1-3) by inserting short RBM sequences in either CDR2 or CDR3 (Fig. 1D-E) to evaluate local folding variability (see Material and Methods for engineering techniques). Model 2 and 3 were also designed to include the sequence QYIKANSKFIGITE, a universal T helper (Th) cell epitope from tetanus toxoid (Panina *et al*, 1989), in CDR3 and CDR2, respectively (Fig. S1). This epitope is not presented by the two classical class II antigens of the H2 complex I-A and I-E, and served to assess possible effects on folding of the B cell epitope and its immunogenicity.

**Fig. 1.**
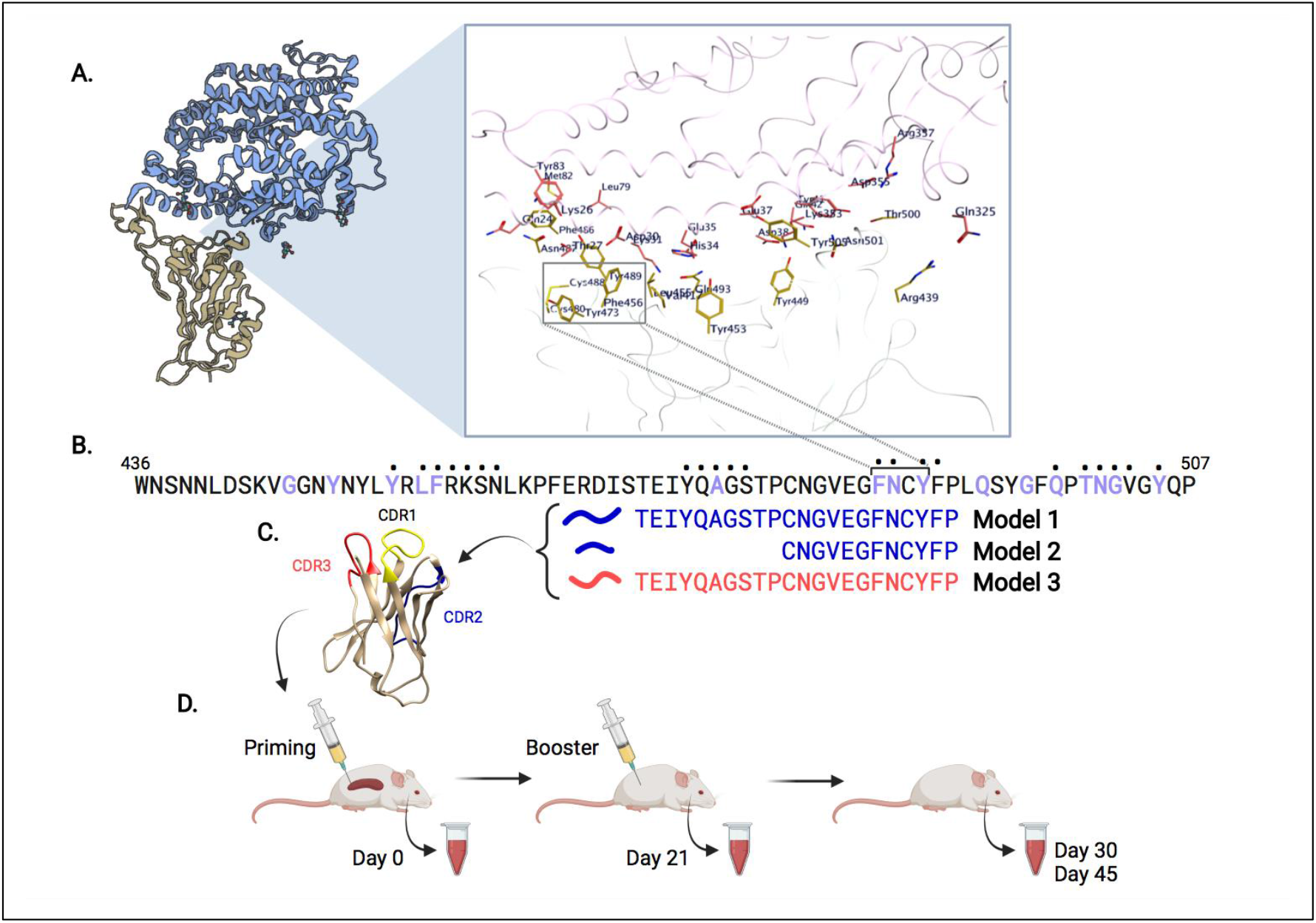
Overview of SARS-CoV-2 epitope selection, protein engineering, and immunization. (**A**) SARS-CoV-2 Spike protein (yellow) interacting with ACE2 (blue), PDB ID: 6M0J. A zoomed view shows SARS-CoV-2:ACE2 interacting residues. (**B**) Spike protein RBM (436-507) sequence. Purple residues indicate ACE2 binding, dots above residues indicate B38 or CC12.1 antibody binding. Immunogens models 1-3 span the putative B cell epitope FNCY (486-489). (**C**) VH62 model with CDR1 (yellow), CDR2 (blue), CDR3 (red). (**D**) Timeline of priming (day 0), and booster shot (day 21), with blood draws (days 0, 21, 30, 45).

All pDNAs are under the control of the Ig promoter so that transcription and translation of the rearranged Ig gene are restricted to B cells. The injection of pDNA into the B cell rich environment of the spleen leads to a process of immunity within the spatio-temporal geometry of an organized secondary lymphoid tissue, mimicking the immunodynamics of viral infection without being infectious with advantages for the formation of long-lasting immunological memory (Zanetti *et al*, 2004).

Female C57BL/6 mice (N=4 per group) were primed by single intra-spleen injection of one of three RBM pDNAs (model 1-3) (Fig. 1E). Mice were given a booster immunization with Spike protein (20 μg) in incomplete Freund’s adjuvant (IFA) on day 21. Group 4 only received the booster immunization with Spike protein in IFA only. Sera were collected before the booster immunization and tested by ELISA on purified RBD and Spike (D614G mutation) proteins. Weak binding to RBD at low serum dilutions (1:50) occurred mainly in group 1 and 2 (Fig. 2B), in line with the characteristics of this form of immunization (Gerloni *et al*, 1997). The response against the Spike protein occurred in group 1 and 2 (Fig. 2C). The stronger binding to the Spike protein likely reflects a better exposure of the RBM since the D614G mutation keeps the protomers in a prevalent “up” position (open conformation) (Benton *et al*, 2020). A primary antibody response was evident in group 1 and 2 suggesting a more favorable surface exposure and spatial orientation of the FNCY patch and surrounding residues in the immunogenic proteins coded by pDNA1 and pDNA2.

**Fig. 2.**
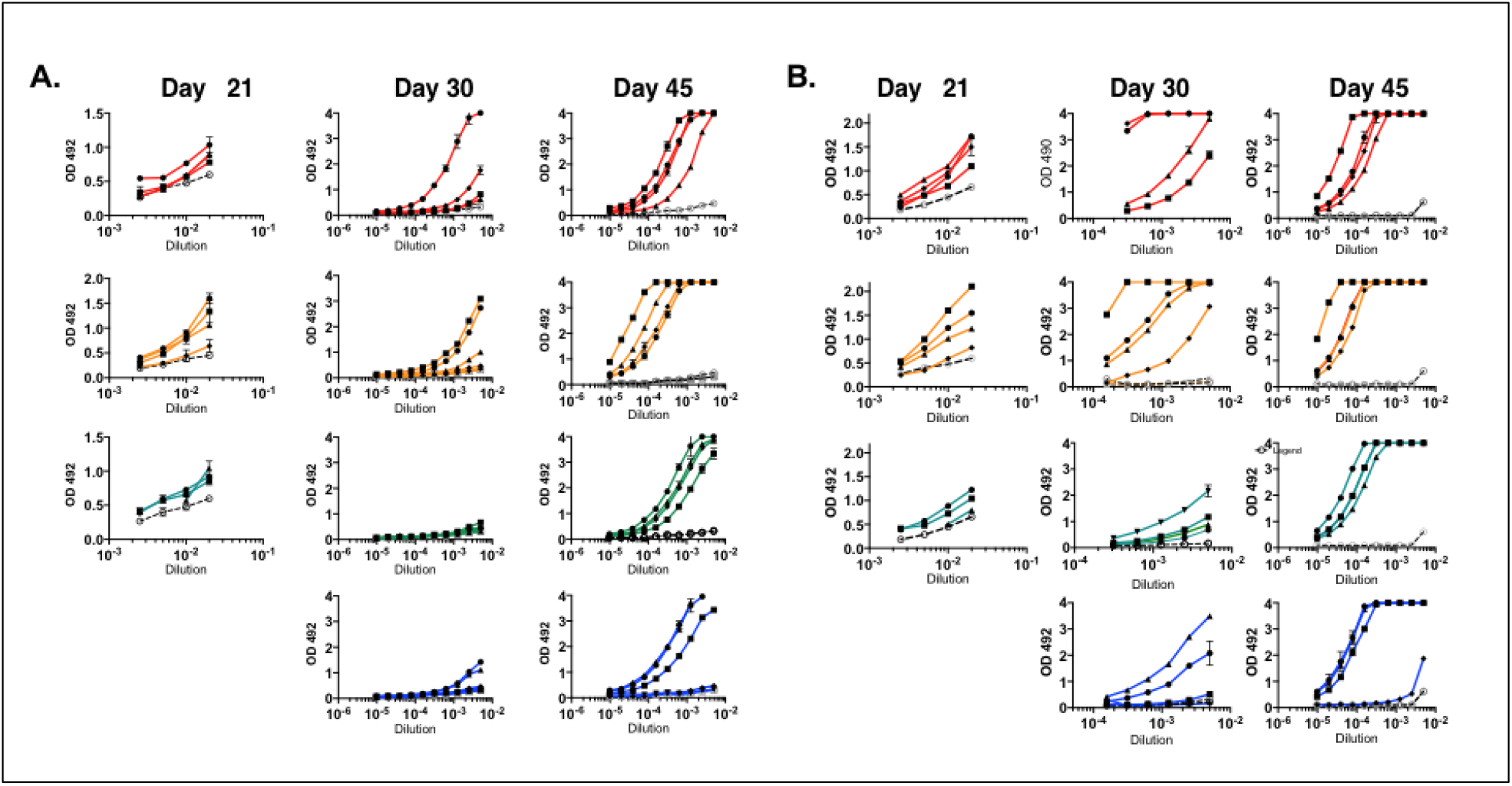
ELISA binding of immune sera of mice after prime-boost. Binding to RBD (**A**) and Spike protein (**B**) of sera at the timepoints indicated. Closed symbols refer to individual mice. Open symbols refer to binding of pre-immune sera. Results are from one experiment representative of three independent runs. Color scheme: red = group 1; orange = group 2; green = group 3; blue = group 4.

The antibody response post-booster immunization was analyzed at an early (day 30) and late (day 45) time point (*i.e*., 9 and 24 days after booster), to capture the evolution of the memory response. Nine days post-boost antibodies to RBD were markedly increased relative to day 21 both in group 1 and 2. The response was also greater than in group 3 and 4. Since group 4 controls for magnitude and speed of the antibody response after booster with intact Spike protein, the data show that pDNA priming accelerated the recall antibody response by the Spike protein. Not surprisingly, sera from group 1 and 2 also had a markedly stronger response to the Spike protein compared to group 3 and 4 (Fig. 2).

Twenty-four days post-boost the responses against the RBD and Spike proteins were considerably stronger than on day 30 and substantially similar among groups (Fig. 2). Collectively, the three pDNA immunogens differed in their ability to prime a specific B cell response and generate an anamnestic response in a prime-boost regimen. A greater response against the RBM B cell epitope was associated with expression of the RBM epitope in the CDR2 loop of the VH, suggesting context-dependent immunogenicity perhaps owing to better folding and more favorable recognition by B cells relative to expression in CDR3.

The B cell epitope selected for these studies is contained in a narrow region of the RBM. To monitor the reactivity of serum antibodies with greater precision we synthesized peptide ^475^AGSTPCNGVEGFNCYFPLQSYGFQPT^500^. Reactivity against the RBM peptide occurred after booster immunization with binding profiles on day 9 post-booster mimicking those on RBD and Spike proteins in group 1 and 2, which displayed overall stronger binding relative to group 3 and 4 (Fig. S2A). Intra-spleen pDNA priming induces predominantly IgM antibodies (Gerloni *et al*, 1998), which may account for a weak binding to the RBM peptide at 1:50 dilution, suggesting low antibody concentration in sera, low avidity, or both. Antibody binding increased by day 45 (Fig. S2A). Thus, priming with pDNA1 and pDNA2 generated a pool of memory B cells specific for an RBM epitope comprised within residues ^475^AGSTPCNGVEGFNCYFPLQSYGFQPT^500^ that were expanded during the anamnestic response.

### Immune sera antibodies cross-compete RBD binding of neutralizing human monoclonal antibodies

The RBM amino acids grafted into CDR loops comprise contact residues shown to be targets of potently neutralizing antibodies derived from COVID-19 patients. Among those are antibodies B38 (Wu *et al*., 2020) and CC12.1 (Rogers *et al*., 2020), which map an overlapping RBM epitope including contact residues Phe - Asn and Tyr of the ^486^FNCY^489^ patch in both cases (Fig 3A).

**Fig. 3.**
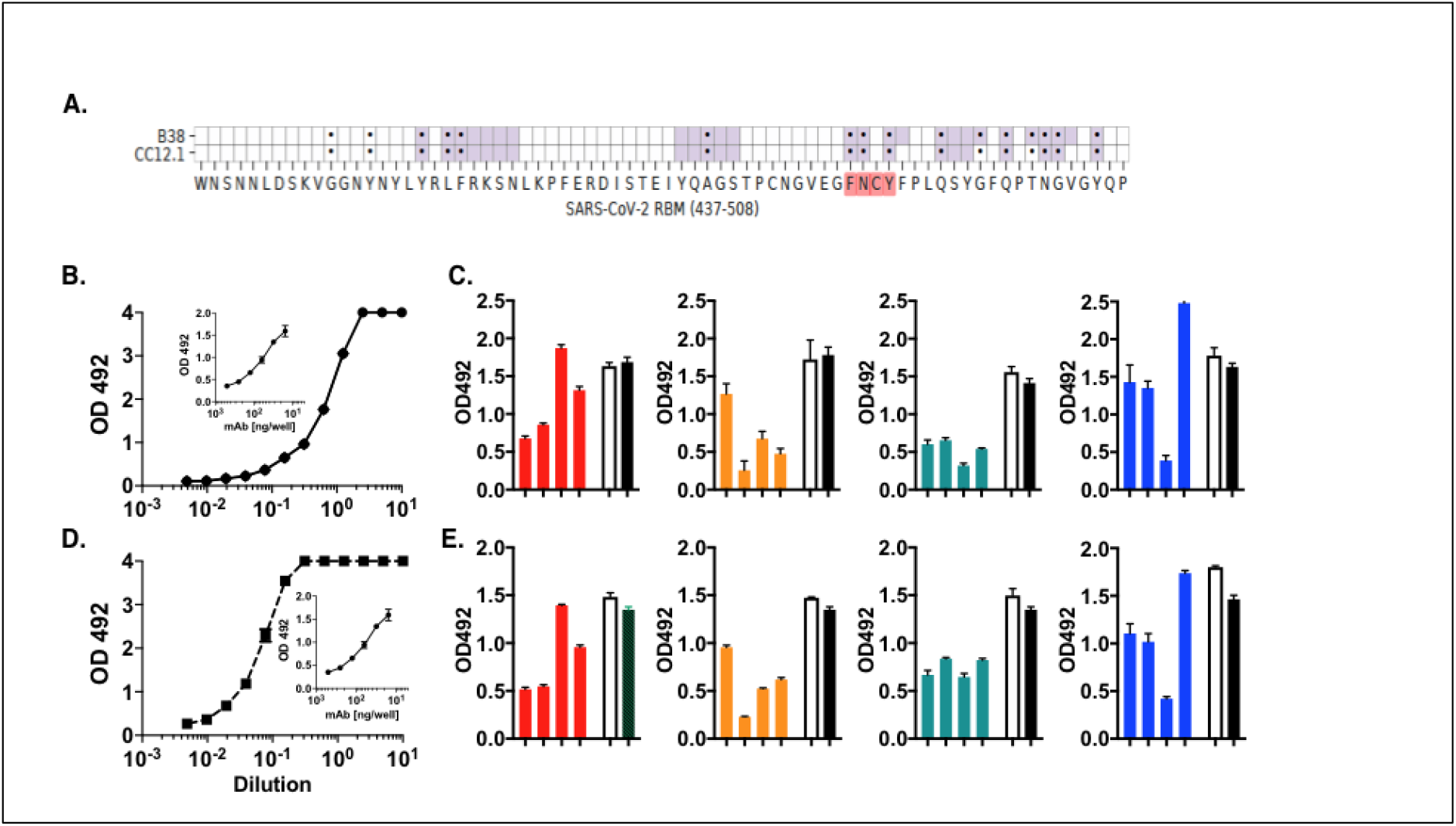
Polyclonal antibodies in immune sera share paratope specificity with Class 1 human neutralizing antibodies B38 and CC12.1. Immune sera were tested in a cross-competition RBD binding assay by ELISA. (**A**) Heatmap of neutralizing antibody contact residues (purple) with RBM region (positions 437−508). Black dots indicate ACE2 contact residues. (**B**) Titration of binding of HRP-B38 to RBD in ELISA. Inset: Slope of inhibition of HRP-B38 binding by unlabeled antibody B38. (**C**) Binding of HRP-B38 in the presence of 1:50 dilution of individual mouse serum (day 45). (**D**) Titration of binding of HRP-CC12.1 to RBD in ELISA. Inset: Slope of inhibition of HRP-CC12.1 binding by unlabeled antibody CC12.1. (**E**) Binding of HRP-CC12.1 in the presence of 1:50 dilution of individual mouse serum (day 45). Empty columns; pre-immune sera. Black columns indicate maximal binding of HRP-labeled antibody in ELISA buffer.

We reasoned that the paratope of these two antibodies could be used to determine shared epitope specificity between immune serum polyclonal antibodies and human neutralizing antibodies. To this end, we designed a competitive ELISA assay where the binding of horseradish peroxidase (HRP)-labeled B38 and CC12.1 to the RBD protein was competed by individual immune sera. HRP-labeled B38 and CC12.1 bound RBD with similar characteristics (Fig. 3B and D) so that both antibodies could be used at the same (~1.5 ng/well) effective ~50% binding concentration. Unlabeled antibodies B38 and CC12.1 inhibited homologous RBD binding at similar concentration (50% inhibition at ~ 60 ng/well) (Fig. 3B and D, inset).

The majority of immune sera tested individually at a 1:50 dilution inhibited >50% the binding to RBD by both HRP-B38 and HRP-CC12.1. Inhibition in group 2 was overall the strongest (range 25-85% for B38; and 30-84% for CC12.1) (Fig 3C and E). Surprisingly, group 3 immune sera also inhibited (range 62-80% on B38; and 38-52% on CC12.1). Group 4 had overall the weakest inhibitory activity (<30%) with the exception of one mouse. The fact that serum antibodies from immune mice cross-competed the RBD binding of the two human neutralizing antibodies indicates that a component of serum antibodies in the immune serum share the paratope with antibodies B38 and CC12.1. We estimated that the upper limit serum concentration of such antibodies as high as ~12 μg/ml. Collectively, the results show that intra-spleen immunization with pDNA coding for the selected RBM epitope activates, and subsequently facilitates, the expansion of B cell clonotypes producing RBM antibodies found in COVID-19 patients. Oddly, neither HRP-B38 nor HRP-CC12.1 bound to ^475^AGSTPCNGVEGFNCYFPLQSYGFQPT^500^ peptide in ELISA (Fig. S2B and C), suggesting perhaps that the synthetic peptide lacks the conformation/structure the paratope of these antibodies needs for binding. Thus, immune sera have a wider spectrum of paratopes than those defined by the two human monoclonal antibodies.

### Neutralization of authentic SARS-CoV-2 isolates WA1 and variants of concern B.1.351 and B.1.617.2

We tested the neutralizing activity of day 45 sera on authentic SARS-CoV-2 isolates, USA-WA1/2020 (WA1) and VOC lineages B.1.351 20H/501Y.V2/Beta (B.1.351) and B.1.617.2 21A/S:478K/Delta (B.1.617.2), in a focus reduction neutralization test (FRNT) (Fig. 4 and Fig. S3). We found that sera from group 1 and 2 gave marked neutralization of WA1 that was titratable over a 3 log_10_ dilution range with IC_50_ +/− SEM of 1:2,588 +/− 993 and 1:1,637 +/− 196, respectively. Group 3 also inhibited WA1 with an IC_50_ of 1:2,321 +/− 756. Reference group 4, which only received the booster immunization of 20ug of spike protein in IFA was neutralizing except in one case. Neutralization by control monoclonal antibodies CC12.1 and CC6.30 was strong (IC_50_ of 116 ng/ml and 8 ng/ml, respectively) consistent with published data. Pre-immune mouse sera did not neutralize. We then assessed neutralization of the B.1.351 VOC.

**Fig. 4.**
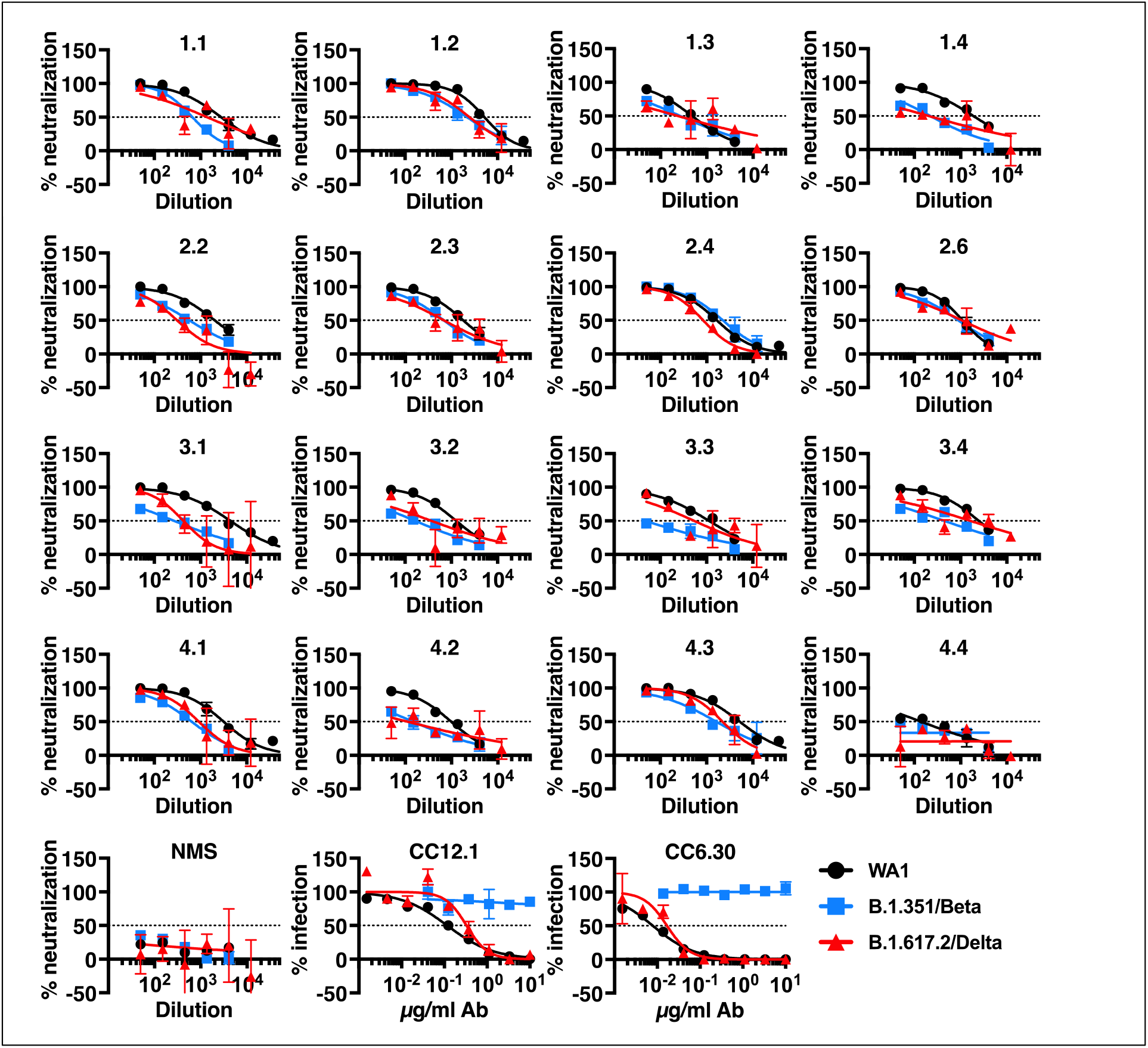
Neutralization of authentic SARS-CoV-2 WA1 isolate and VOCs B.1.351/Beta and B.1.617.2/Delta. Neutralization of authentic SARS-CoV-2 isolate WA1 and VOCs B.1.351 and B.1.617.2 by immune sera was measured by focus reduction neutralization test (FRNT) on TMPRSS2-Vero cells. 3fold serial dilutions started at 1:50. Percent neutralization and % infection are relative to media-only control. Data shown are the mean and SEM of duplicate virus + serum incubations. Dotted lines mark 50% neutralization. Representative ot two independent runs.

The B.1.351 variant has multiple mutations in the spike protein, including the K417N, which is outside the RBD, and the E484K and N501Y mutations that are within, or immediately adjacent to the RBM ridgeline. These mutations are also present in SARS-CoV-2 VOC P.1 and some isolates of B.1.1.7, and contribute to reduced neutralization by monoclonal antibodies, as well as convalescent and post-vaccination sera (Garcia-Beltran *et al*, 2021; Wang *et al*, 2021b; Weisblum *et al*, 2020). We noted that group 2 had consistent neutralization of B.1.351. In one instance, neutralization of both the WA1 and B.1.351 isolates was nearly superimposable. Group 1 sera also neutralized B.1.351 infection, but less consistently compared to wild type virus. In contrast, neutralization of the B.1.351 variant by group 3 sera was poor or absent. Group 4 showed cross-neutralization in 2 out of 4 instances. Neither monoclonal antibody CC12.1 nor CC6.30 neutralized the B.1.351 variant. Together, these results suggest that pDNA priming with the RBM epitope in CDR2 synchronizes a response to subsequent booster protein immunization that privileges recognition of both wild type SARS-CoV-2 and the B.1.351 VOC. Abrogation of neutralization of the B.1.351 VOC by monoclonal antibodies CC12.1 and CC6.30 is consistent with the key role of mutation K417N in abrogating the binding af neutralizing antibodies belonging to the Class 1 group (Yuan *et al*, 2021).

The B.1.617.2 variant has several mutations in the spike protein, including L452R and T478K (two non ACE2 contact residues) and P681R in the S2 subunit, but not the K417N, E484K and N501Y mutations present in the B.1.351 VOC. While again neutralization occurred in all groups we noted that group 2 had consistent neutralization of B.1.617.2 in much the same way as WA1 and B.1.351. In other groups, IC_50_ for B.1.617.2 was intermediate between values for WA1 and B.1.351, and the monoclonal antibodies CC12.1 and CC6.30 neutralized effectively as predicated by the lack of K417N mutation.

In conclusion, we demonstrate that immune mice sera neutralize not only authentic WA1 virus but also two VOCs responsible for rapid spreading of infection and disease.

### Structure-function considerations on B cell epitope expression

We used computer modeling techniques to model the RBM B cell epitope CNGVEGFNCYFP expressed in the CDR2 of Model 2 as mice primed with pDNA 2 provided the most consistent antibody response and neutralized consistently both authentic wild type virus and the B.1.351 and B.1.617.2 VOCs. Model 2 expresses RBM epitope in the CDR2 region and the sequence QYIKANSKFIGITE, a universal T helper epitope from the tetanus toxoid (TT) in CDR3. However, the TT epitope is not presented by the two classical class II antigens of the H2 complex, I-A and I-E. Therefore, its presence is only relevant to folding and immunogenicity of the RBM B cell epitope.

As seen in Fig. 5B and C, the orientation of the RBM epitope, notably the FNCY patch, is projecting outward and is solvent-exposed as in the Spike 1 protein. Examining the docking interactions between the Phe-Asn-Tyr triad and ACE2 residues (Fig. 5D and E; Table S1) it appears that the RBM epitope expressed in the CDR2 loop establishes a greater number of shorter distance binding interactions with ACE2, suggesting an overall stereochemical similarity with the corresponding FNCY patch residues in the virus RBM.

**Fig. 5.**
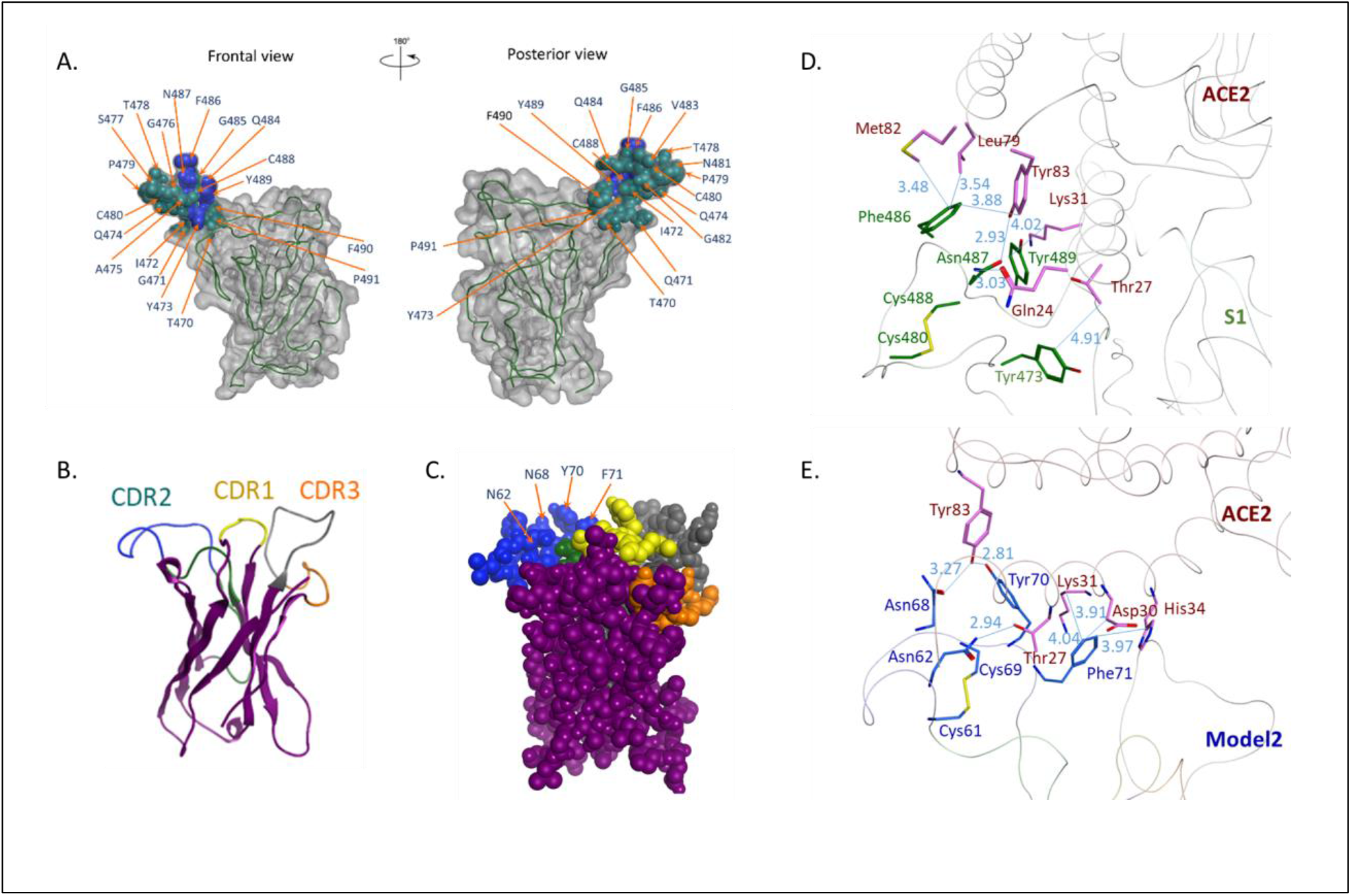
Conformational analysis of SARS2-CoV-2 RBM structure and predicted conformation of VH Model 2. In the Spike S1 protein (**A**) the tip of the RBM ridgeline is shown in dark green and the FNCY patch in dark blue. In Model 2 (ribbon **B**) and (space filling **C**) the modeled CDR2 loop is shown in dark green, while the grafted RBM epitope is in dark blue. Panels on the right show the interface between ACE2 and SARS-CoV-2 RBM (**D**), and ACE2 and the CDR2 loop of Model 2 (**E**). Color scheme: A: gray - RBD; green - RBM; blue – ACE2 contact residues; B and C: yellow - CDR1; green - CDR2; orange - CDR3; blue - SARS-Cov-2 RBM residues grafted in CDR2; gray - tetanus toxoid epitope; D: pink - ACE2; green - SARS-Cov-2 RBM residues; E: pink - ACE2; blue - SARS-Cov-2 RBM residues grafted in CDR2.

## Discussion

We engineered three immunogens expressing a B cell epitope of the RBM ridgeline in the CDR loops of an Ig V domain scaffold as immunogens against SARS-CoV-2. The B cell epitope spans a small region (22 amino acids) of the RBM ridgeline and encompasses the solvent-exposed ^486^FNCY^489^ patch, which contributes three contact residues to ACE2 binding (Lan *et al*., 2020). Numerous human Class 1 neutralizing antibodies (Barnes *et al*., 2020a), including B38 and CC12.1, have been mapped to this region (Rogers *et al*., 2020; Wu *et al*., 2020). We used a DNA-protein (heterologous prime-boost) approach to screen pDNAs able to prime an antibody response against RBD and induce a memory response. We show that two of the three pDNAs tested, both expressing the RBM B cell epitope in CDR2, induced a strong neutralizing response not only against the WA1 isolate but also the B.1.351 and B.1.617.2 VOCs.

After pDNA priming, polyclonal serum antibodies (group 1 and 2) bound the RBD and the Spike proteins, and also synthetic peptide ^475^AGSTPCNGVEGFNCYFPLQSYGFQPT^500^. Interestingly, ELISA titers at the early (day 30) expansion of the memory response correlated with a consistent virus neutralization on day 45 (group 2), suggesting that the expansion of B cells specific for the minimal RBM B cell epitope ^480^CNGVEGFNCYFP^491^ by pDNA priming generated memory B cells that were readily expanded by booster immunization. Analysis of day 45 immune sera showed strong binding to RBD and Spike proteins, but also inhibition of binding to RBD by two potently neutralizing human antibodies (B38 and CC12.1) reported to map to an overlapping site in the RBM ridgeline (Fig. 3A). This suggests that some among the polyclonal antibodies in immune sera shared epitope specificity with these two human neutralizing antibodies. Moreover, since neither antibody B38 nor CC12.1 bound peptide ^475^AGSTPCNGVEGFNCYFPLQSYGFQPT^500^, it appears as if polyclonal antibodies have multiple paratope specificities and that unlike antibodies B38 and CC12.1 their binding to the RBM B cell epitope is independent of residue K417 (Yuan *et al*., 2021). An alternative explanation is that their paratopes bind from a different angle.

Polyclonal antibodies in immune sera after prime-boost immunization neutralized efficiently authentic SARS-CoV-2 WA1, B.1.351, and B.1.617.2 isolates. Although neutralization of wild type virus was expected in light of the strong ELISA antibody titers against RBD and Spike proteins, neutralization of the B.1.351 and B.1.617.2 VOCs, particularly by group 2 sera, was not. Compellingly, this shows that few critical solvent-exposed amino acids in the RBM can expand B cells with a paratope specific for the RBM. One may argue that neutralization of the two VOCs by polyclonal antibodies in immune sera is owed to recognition of a discrete epitope within the RBM that is conserved among the WA1 virus and its two variants B.1.351 and B.1.617.2. In line with this interpretation Wang et al. (Wang *et al*, 2021a) showed that potent neutralizing antibodies in convalescent individuals that utilize the IGVH1.58 germline gene have similar potency against the B.1.351 and B.1.617.2 variants. These antibodies map to a “supersite” that is overlapping with the minimal RBM B cell epitope ^480^CNGVEGFNCYFP^491^ characterized in the present study. Collectively, this shows that when this B cell epitope is conformationally-constrained in the CDR2 is not only immunogenic but also captures a spectrum of paratopes efficient at neutralizing across the WA1 virus, and the B.1.351 and B.1.617.2 variants of concern. This stresses the importance of defining immunogenic sites at the structure/function level for efficient precision immunization.

Our study has limitations. First, the sample size was deliberately low since the study was designed to identify (a) pDNA with ability to prime RBM-specific B cells that could be re-expanded during a recall response, and (b) yield consistent virus neutralization. In spite of this limitation we managed to identify the immunogen yielding the overall more consistent response. The second is that the dose of Spike protein used in the booster immunization was probably excessive as this protein is highly immunogenic. This may have masked the true potential of the memory response. Future experiments will need to assess memory responses against booster immunization with lower doses of antigen. Finally, although the study was successful in identifying an optimal prime-boost combination resulting in neutralizing antibodies effective against wildtype virus and B.1.351 and B.1.617.2 variants, these results cannot be extrapolated to predict transmission inhibition *in vivo*.

Control of the COVID-19 pandemic rests on an effective immunological intervention to curb the spread of infection by vaccination to induce durable transmission-blocking immunity. The initial evidence gathered in this study suggests that a pDNA-protein (prime-boost) approach was successful in focusing the antibody response to a narrow site of the RBM ridgeline that overlaps with the RBM supersite recently described (Wang *et al*., 2021a) and proved effective against wildtype virus and two VOCs. This suggests that binding and neutralization mediated by polyclonal antibodies in immune sera is likely enriched in antibodies binding the RBM supersite. Our data also show that the E484K mutation which is present in the B.1.351 VOC and shared by another VOC (P.1 or 20J/501Y.V3) as well as several variants of interest (B.1.525, P.2, P.3, and some isolates of B.1.526) does not block the reactivity of antibodies generated in our prime-boost immunization. This contrasts studies showing that antibodies induced by natural infection or vaccines based on wildtype Spike are less effective at neutralizing the B.1.351 variant (Cele *et al*, 2021; Collier *et al*, 2021; Jangra *et al*, 2021; Liu *et al*, 2021; Shen *et al*, 2021; Wang *et al*., 2021b; Wu *et al*, 2021), leading potentially to immune escape.

The results presented here show that immunogens expressing a preselected site of the RBM ridgeline can focus the antibody response to the RBM. The need to do so is emphasized by the fact that > 80% of the whole antibody response in humans is directed predominantly to sites outside the RBD (Voss *et al*., 2021) consistent with the observation that B and T cell responses targeting the RBD, and in particular the RBM, are markedly less frequent than the total response to the S protein (Juno *et al*, 2020). However, all but one of 20 most potent neutralizing antibodies (IC_50_ < 0.1 μg/mL) characterized to date bind the RBM and block receptor attachment (Dejnirattisai *et al*., 2021).

The type of immunogens tested here were designed to prime the antibody response against a selected site in the RBM ridgeline which appears to overlap with the RBM supersite recognized by neutralizing antibodies utilizing the IGVH1.58 gene (Wang *et al*., 2021a). They also test the hypothesis that protective B cell immunity can be initiated within the spatial structure and community of resident immune cells in a secondary lymphoid organ. Therefore, it will be important to develop delivery strategies that mimic the approach demonstrated here, i.e., initiation of immunity at the site of immune induction. While our modular engineering approach can be easily expanded to express in conjunction additional B cell epitopes mapping to sites of vulnerability of the virus, we propose such an approach to also limit the expansion of B cells clones selected on immunodominance (Altman *et al*., 2018) that may reactivate the production of pathogenetic autoantibodies (Bastard *et al*, 2020; Zuniga *et al*, 2021; Zuo *et al*, 2020), and cause immune suppression (Xu *et al*., 2018).

Achieving global control of this pandemic will require vaccines that overcome obstacles such as antigen stability, vaccine thermostability, and the logistics of cold chain requirements (Agrawal *et al;* Capua & Giaquinto, 2021). pDNA vaccines of the type presented here offer such a possibility; for instance they can be incorporated in thermostable needle-free delivery vehicles for global and equitable vaccination (Nachega *et al*, 2021).

## Materials and Methods

### Experimental Design

The experimental design is illustrated in Fig. 1. The overall objective of the study was to identify an immunogen able to prime a polyclonal response expandable as a memory response upon boost immunization while enriching for antibodies recognizing a specific site in the RBM ridgeline.

### Recombinant proteins, monoclonal antibodies and synthetic peptide

The generation of SARS-CoV-2 HexaPro Spike variant D614G and soluble RBD (sRBD) was performed as follows. SARS-CoV-2 HexaPro ectodomain containing residues 14-1208 (Genbank: MN908947) with the mutation D614G was stabilized in the prefusion conformation by the introduction of six proline substitutions (F817P, A892P, A899P, A942P, K986P, V987P), the replacement of cleavage site residues 682-685 (“RRAR” to “GSAS”), and the introduction of a C-terminal Foldon trimerization domain. For the generation of a soluble version of the SARS-CoV-2 RBD, one gene encoding amino acids 319-591 from the Wuhan variant was used. Both proteins HexaPro and sRBD were then cloned into a phCMV mammalian expression vector containing an N-terminal Gaussia luciferase signal sequence (MGVKVLFALICIAVAEA) and two C-terminal Strep-Tags for the purification of the proteins. Between the proteins and the purification tags an HRV-3C cleavage site was introduced to enzymatically remove the tags after the purification if necessary. Plasmids were transformed into Stellar competent cells and isolated using a Plasmid Plus Midi kit (Qiagen). Transient transfection and protein purification were performed as follows. Briefly, SARS-CoV-2 Spike ectodomain and the sRBD were transiently transfected in Freestyle ExpiCHO-S cells (Thermo Fisher). ExpiCHO cells were maintained and transfected according to the manufacturer using the “High Titer” protocol. Briefly, plasmid DNA and Expifectamine were mixed in Opti-PRO SFM (Gibco) according to the manufacturer’s instructions, and added to the cells. One day after the transfection, cells were fed with manufacturer-supplied feed and enhancer as specified in the manufacturer’s according to protocol, and cultures were set to 32 °C, 5% CO_2_ and 115 RPM. One week later, supernatants were clarified by centrifugation, BioLock was added, passaging through a 0.22 μuM sterile filter, and proteins were first purified on an ÄKTA go system (Cytivia) using a 5mL StrepTrap-HP column equilibrated with TBS buffer (25mM Tris pH 7.6, 200mM NaCl, 0.02% NaN33) and eluted using TBS buffer supplemented with 5mM d-dDesthiobiotin (Sigma Aldrich). Proteins were then second purified by size-exclusion-chromatography (SEC) on a Superdex 6 increase 10/300 column (Cytivia) in the same TBS buffer. Human monoclonal antibodies B38, CC12.1 and CC6.30 have been described previously (Rogers *et al*., 2020; Wu *et al*., 2020). The RBM peptide ^475^AGSTPCNGVEGFNCYFPLQSYGFQPT^500^ was synthesized by Synthetic Biomolecules (San Diego) and purified by HPLC (>90% purity).

### Protein Engineering methods

Engineering methods were a modification of those described in (Xiong *et al*, 1997). Briefly, the three VH genes matching the descriptions of Model 1-3 as shown in Fig. 1 were synthesized with unique ends for cloning into the ZUC1.1 (9.7 Kb) target vector using the cloning site as shown in Fig. S4. The ZUC1.1 plasmid lacks Amp resistance gene and SV40 sequences, and is optimized for human use. The amino acid sequences of the Variable domain of Model 1-3 are shown in Fig. S1. pDNAs were prepared from transformed DH5a *Escherichia coli* according to standard procedures and were analyzed for purity using the following equation: %N=(11.1R-6.32)/(2.16-R) where R=260nm/ 280nm, %N=% of nucleic acid ^42^. pDNAs were resuspended in sterile saline solution and stored at - 20 °C until use.

### Mice and immunizations

Twelve-week-old female C57Bl/6 (H-2^b^) were bred at the University of California, San Diego animal facility where they were kept throughout the performance of the experiment. Mice were primed by single intra-spleen inoculation of 100 μg of plasmid DNA in 50 μl of sterile saline solution. Booster immunization was administered on day 21 after priming by a 2-3 subcutaneous injections of purified Spike protein (20 μg per mouse) emulsified in incomplete Freunds’ adjuvant (IFA). Procedures were per protocol approved by the Institutional Animal Care and Use Committee (IACUC) and in compliance with Association for Assessment Accreditation of Laboratory Animal Care (AAALAC) International guidelines.

### Serologic assays

<u>Direct ELISA</u>. Antibodies to Spike, RBD and RBM peptide were detected by ELISA on 96-well polyvinyl microtiter plate (Dynatech, Gentilly, VA) coated with Spike or RBD (4 μg/ml) or RBM peptide (6 μg/ml) in carbonate buffer pH 8.6, 0.1M, by incubation overnight at +4 °C. After coating, wells were blocked with 5% BSA in PBS for 1 hour at 37 C. Mouse sera were diluted in phosphate buffered saline (PBS) 0.15 M, pH 7.3, containing in 1% bovine serum albumin (BSA) and 0. 5 % Tween-20 and then incubated overnight at +4 °C. The bound antibodies were revealed using a HRP-conjugated goat antibody to mouse Ig absorbed with human Ig (Sigma) (1:4,000 dilution). The bound peroxidase was revealed by adding o-phenylenediamine dihydrochloride and H_2_O_2_. Plates were read after 30 minutes in a micro-plate reader (TECAN Spark plate reader) at 492 nm. Tests were run in duplicate and repeated 2-3 time for consistency. <u>Competition ELISA</u>. (1) The detection of antibodies in immunized sera with shared epitope recognition with human monoclonal antibodies B38 and CC12.1 was performed as follows. Briefly, B38 (10 μg) and CC12.1 (100 μg) were coupled with HRP using the ab 102890 – HRP Conjugation kit (Lightning Link, Abcam) following the manufacturer’s instructions. Sera from individual mice (1:100 dilution or otherwise specified) in PBS-BSA containing 0.5 % Tween-20, were incubated overnight with a dilution of HRP-B38 or HRP-CC12. 1 (~2.5 ng/ml) at +4 °C on 96-well plates coated with RBD protein (4 μg/ml). Binding was revealed as described above.

### SARS-CoV-2 viruses

All work with SARS-CoV-2 was conducted in Biosafety Level-3 conditions at the University of California San Diego following the guidelines approved by the Institutional Biosafety Committee. SARS-CoV-2 isolates WA1(USA-WA1/2020, NR-52281) and B.1.351/South Africa/Beta (hCoV-19/South Africa/KRISP-K005325/2020, NR-54009) were acquired from BEI and passaged once through primary human bronchial epithelial cells (NHBECs) differentiated at air-liquid interface (ALI) to select against furin site mutations. Culture and differentiation of NHBECs at ALI are described below. Viruses were then expanded by one passage through TMPRSS2-Vero cells. Isolate B.1.617.2 (hCoV-19/USA/PHC658/2021, NR-55611) was expanded on TMPRSS2-Vero cells. Supernatants were clarified and stored at - 80°C and titers were determined by fluorescent focus assay on TMPRSS2-Vero cells. Sequence of viral stock was monitored by RNAseq.

### Primary Normal Human Airway Epithelial Cell Culture and Differentiation at Air-Liquid Interface

Primary normal human bronchial epithelial cells (NHBECs) were obtained from a 65 year old Caucasian male without identifiers sourced from Lonza (NHBE CC-2540; Walkersville, MD). The NHBECs were revived from cryopreservation and expanded with PneumaCult™ Ex-Plus media (StemCell #05040, Tukwila, WA). 50,000 cells were seeded on transwells (Corning #29442-082, VWR) pre-coated with 50ug/mL Collagen type I from Rat tail (BD Biosciences #354236) at 7.5ug/cm2 in PneumaCult ^™^ Ex-Plus media. Media was changed on days 1 and 3. On Day 4-7, the apical and basolateral chambers were fed PneumaCult ^™^ ALI media (StemCell #05021) supplemented with 10μM ROCK inhibitor (Tocris Y-27632). Upon reaching confluency, the apical media was removed (airlift) and the basal media replaced with PneumaCult ^™^ ALI media without Y-27632. Subsequent media changes were every 2-3 days thereafter. On Day 14 post-airlift, the apical surfaces were washed with DPBS once per week. Cells were grown in 37°C, 5% CO_2_ incubator until four weeks airlifted.

### Infection of NHBECs at air liquid interface

After two 30 min incubations with PBS at 37°C, 5% CO_2_, virus diluted in PBS was added to apical chambers in 100uL. Virus was removed after 24h and apical washes (150uL PBS with 10 min incubation at 37°C, 5% CO_2_,) were taken daily and stored at −80. Titers were determined by fluorescent focus assay on TMPRSS2-Vero cells.

### Authentic virus neutralization assay

Neutralizing antibodies in immune sera were screened through the focus reduction neutralization test (FRNT). Briefly, ~100 focus forming units (ffu) of SARS-CoV-2 were incubated with or without serially diluted antibodies for 1 hour in DMEM with 1% FBS at 37 °C before adding to 100% confluent TMPRSS2-Vero cell monolayers in 96 well plates. After 1 hour incubation at 37 °C with rocking, inocula were removed and monolayers overlaid with 100μl of DMEM + 2% FBS containing 1% methylcellulose. After 1 day incubation (37 °C) cells were fixed with 4% formaldehyde and stained with antibody against nucleocapsid protein. Plates were imaged on an Incucyte S3 and foci were counted. Neutralization was calculated as the percentage of reduced foci compared to the mean of the foci in virus media-only control wells on each plate. Best fit curves generated in PRISM 9 determined the serum dilution which achieved 50% focus reduction (FRNT_50_).

### Computational modeling

Computational models utilized the coordinates of rearranged murine VH62 (Sollazzo *et al*, 1989). Model predictions have made using SWISS-MODEL, an automated protein structure homology-modelling server and the coordinates of the crystal structure of an antibody: antigen complex (SMTL ID: 5mhe.1) where the antibody is specific for L-Thyroxine, which is also the epitope specificity of monoclonal antibody 62 from which VH62 was originally cloned (Sollazzo *et al*., 1989). Structure predictions of the SARS-Cov2 RBM B-cell epitope CNGVEGFNCYFP inserted in CDR2 were made using Loop Modeler application of Molecular Operating Environment (MOE) program (Chemical Computing Group, Montreal, Canada). A set of variants were subsequently generated. Selected conformation of Model 2 antibody was saved. The CDR3 was modified by insertion of the universal T helper (Th) cell epitope from tetanus toxoid QYIKANSKFIGITE and was modified using Loop Modeler of MOE. The CDR1 was not modified after SWISS-MODEL. Then the entire antibody model was docked to the ACE2 protein, using Dock module of MOE software and the interface analyzed to elucidate the interacting residues of both proteins.

### Quantification of data and statistical analysis

Data plotting and statistical analyses were done using GraphPad Prism version 7.0. Data plotted in logarithmic scales are expressed as Mean + Standard Deviation (SD). Additional data analysis techniques are described in the Methods sections above.

## Acknowledgments

The authors thank Dr. Dennis Burton (The Scripps Research Institute, La Jolla, CA) for the generous gift of purified human monoclonal antibodies CC12.1 and CC6.30. The following reagent was deposited by the Centers for Disease Control and Prevention and obtained through BEI Resources, NIAID, NIH: SARS-Related Coronavirus 2, Isolate USA-WA1/2020, NR-52281. The following reagent was obtained through BEI Resources, NIAID, NIH: SARS-Related Coronavirus 2, Isolate hCoV-19/South Africa/KRISP-K005325/2020, NR-54009, contributed by Alex Sigal and Tulio de Oliveira. The following reagent was obtained through BEI Resources, NIAID, NIH: SARS-Related Coronavirus 2, Isolate hCoV-19/USA/PHC658/2021 (Lineage B.1.617.2; Delta Variant), NR-55611, contributed by Dr. Richard Webby and Dr. Anami Patel.

## Author contributions

Conceptualization: MZ

Methodology: GA, VK, AC, CW, JRK

Investigation: GA, VK, AC, EO, YW

Visualization: VA, AC

Supervision: IFT, GFG, DRB, EOS

Writing—original draft: MZ, AFC, VK, IFT

Writing—review & editing: MZ, AFC, VK

## Data and materials availability

All data are available in the main text or in the Supplementary Materials.

